# Single-cell-based evidence for GLS1 inhibitor as a bona fide senolytic agent in vivo

**DOI:** 10.1101/2025.07.22.666220

**Authors:** Yuki T. Okamura, Teh-Wei Wang, Satoshi Kawakami, Taiki Morimura, Yoshikazu Johmura, Makoto Nakanishi

## Abstract

A plethora of senolytic compounds and technologies have been identified. However, the efficacy of these treatments in eliminating senescent cells and the suppression of chronic inflammation remains to be substantiated in vivo. Here, we employ single-cell RNA sequencing and find the selective elimination of highly inflammatory fibroblasts, endothelial cells in the lung, and proximal tubule cells in the kidney from the GLS1 inhibitor BPTES-treated aged mice. The eliminated fibroblasts share typical phenotypes with in vitro human senescent cells. These cells predominantly express Dpp4 (CD26) and Cadm3, showing activated IFN signaling and lysosomal membrane damage. Cells eliminated by BPTES in lungs exhibit transcriptomes similar to those eliminated in p16-DTR mice, where DTR expression is limited in p16-expressing cells. BPTES treatment results in the T cell population shift from cytotoxic to a protective state, suggesting the suppression of age-related chronic inflammation. CellChat analysis revealed that multiple cytokine signals are transmitted from inflammatory fibroblasts and proximal tubular cells to immune cells in lungs and kidneys. These results provide evidence that GLS1 inhibitor functions as a bona fide senolytic drug to eliminate inflammatory cells, including a subset of senescent cells, and suppresses age-related chronic inflammation.

## Introduction

Aging is an inevitable phenomenon in human life, characterized by a decline in the function of all organs and tissues. Conventionally, the aging process has been regarded as a random phenomenon of life occurring in accordance with the second law of thermodynamics [1]. However, recent studies on the aging process in other species have demonstrated that certain organisms exhibit minimal or no aging, suggesting that aging may not be an inevitable phenomenon of life [2–5]. Aging is a major causative factor in numerous human diseases, including neurodegenerative diseases and cancer. One of the primary pathological phenomena associated with the aging process is “chronic inflammation,” which significantly contributes to the onset of many diseases associated with organ and tissue dysfunction and the aging process [6,7]. In humans, widespread chronic inflammation has been observed in various organs and tissues of the elderly, with its severity increasing with age [8]. A salient observation is the propensity of long-lived individuals to manifest diminished levels of chronic inflammation. This finding aligns with the established principle that aging phenomena are also observed in organisms with minimal aging processes [9].

The mechanisms underlying the induction of chronic inflammation with aging remain poorly understood. However, recent studies have indicated a potential involvement of cellular senescence in this process [10]. It has been observed that senescent cells secrete various inflammatory cytokines, a phenomenon referred to as the “senescence-associated secretory phenomenon (SASP),” which results in the induction of chronic inflammation [11]. In the 2010s, Baker et al. developed a model using genetic engineering to induce apoptosis in senescent cells in a drug-dependent manner [12,13]. Their findings demonstrated that the artificial removal of p16-high cells from aged individuals suppressed the onset of age-related diseases, including cancer, kidney dysfunction, dementia, and extended a healthy lifespan. This finding suggests that the elimination of senescent cells effectively suppresses the aging process and the onset of age-related diseases. Consequently, the development of technology for the removal of senescent cells has emerged as a topic of global interest.

In accordance with the aforementioned concept, a considerable number of senolytic drugs targeting various molecules and pathways have been identified. A substantial body of research has demonstrated that these compounds can elicit advantageous effects in aged subjects and in disease model mice. The SCAPs (Senescent Cell Anti-Apoptotic Pathways) represent an apoptosis-inhibitory cascade in senescent cells, and 46 molecules targeting SCAPs have been identified as potential senolytic candidates [14]. Dasatinib (D), a multi-receptor tyrosine kinase (RTK) inhibitor; quercetin (Q), a flavonoid polyphenol; and ABT-263, a Bcl-2 inhibitor, were identified as first-generation senolytics [15,16]. D+Q therapy has been scientifically validated for its efficacy in addressing a variety of health issues, including but not limited to frailty [14,17], osteoporosis [18], liver cirrhosis [19], insulin resistance [20], neurodegenerative diseases [21], motor function [17], pulmonary fibrosis [22], and chronic kidney disease [23]. Furthermore, this therapeutic modality has been demonstrated to be associated with an augmented lifespan [14,17]. We also previously identified a GLS1 inhibitor, BPTES, as a senolytic drug selectively targeting cells with lysosomal membrane damages [24]. GLS1 inhibitors have been reported to ameliorate the various age-related disorders [24–27]. However, the efficacy of these senolytic drugs in removing highly inflammatory senescent cell populations in vivo remains to be elucidated. Additionally, the potential of senescent cell removal to ameliorate age-related chronic inflammation is not yet fully understood. Consequently, the in vivo Proof of Concept (POC) for senolytic drugs remains to be established, a deficiency that may act as a substantial impediment to their clinical application.

In order to address these important issues and obtain robust evidence of BPTES, a GLS1 inhibitor, as a senolytic drug in vivo, we performed single-cell RNA-seq (scRNAseq) analysis of lung and kidney from mice with or without treatment with BPTES to determine which cells are affected by BPTES in vivo. We found that BPTES treatment indeed eliminated highly inflammatory cell clusters in both pulmonary fibroblasts, endothelial cells and kidney proximal tubular cells. Pulmonary fibroblasts eliminated by BPTES shared various senescence hallmarks with human senescent HCA2 fibroblasts. Importantly, BPTES treatment resulted in a significant suppression of chronic inflammation. Thus, we obtained robust evidence of in vivo POC of BPTES as a senolytic drug in vivo.

## Results

### BPTES eliminated highly inflammatory pulmonary fibroblasts in aged mice

A considerable number of senolytic drugs have been identified, exhibiting the capacity to selectively induce cell death in senescent cells in vitro. A number of these pharmaceutical agents have been documented to alleviate specific age-related dysfunction and diseases. However, there is a paucity of evidence supporting the efficacy of these drugs in eliminating highly inflammatory cells, such as senescent cells, in vivo, and in ameliorating age-related chronic inflammation. These findings are critically important for the clinical application of senolytic drugs in treating age-related diseases. It has been demonstrated that BPTES possesses the capacity to eliminate senescent cells in vitro and to ameliorate various age-related dysfunction and diseases [24]. The underlying mechanism pertains to the failure of neutralization of intracellular acidosis by suppression of ammonia production via GLS1 catalysis from glutamine to glutamate. A subset of senescent cells exhibited evidence of lysosomal membrane damage, which, in turn, triggered an intracellular acidification process.

Therefore, the initial objective was to obtain robust evidence that GLS1 inhibitor selectively eliminate highly inflammatory non-immune cells, such as senescent cells, thereby suppressing age-related chronic inflammation. To this end, we conducted scRNA-seq analysis of lung and kidney samples from aged (20-month-old) mice with or without BPTES treatment (Fig. 1A). The transcriptomes of the entire lung cells were annotated as a total of 32 different cell types (Fig. 1B). Basically, the entire picture of UMAP clustering was very similar to each other with a slight difference in fibroblasts. Then, fibroblasts were subset and reclustered more in detail. Intriguingly, the population of cluster 9 (C9) was drastically reduced in BPTES-treated fibroblasts compared to mock control (Fig. 1C, D). Other clusters (cluster 1 and 2) were also reduced. Cells in C9 exhibited upregulation of known senescent markers, including Cdkn1a and Glb1, and inflammatory genes, such as Il33 (Fig. 1E). More importantly, Gls expression in C9 was significantly higher than other clusters (p.adj=8.8e-14). We further re-analyzed the consistency of transcriptomic alterations between human senescent foreskin fibroblasts induced by nutlin-3a (nSen HCA2 cells)[28] and cells in cluster 9 (Fig. 1F). nSen HCA2 cells were previously proved which sensitized to BPTES-dependent senolysis. We found that 41 genes within C9 specifically expressed genes (274 genes) were overlapped with upregulated genes in nSen HCA2 cells. These overlapped genes included Cxcl1, Ccl2, Il33, Dpp4 (CD26), Cadm3, Wnt5a, and Sfrp2. Among these, Dpp4 and Cadm3 are cell-surface markers that can be used to isolate cells with their specific antibodies [29]. Indeed, our previous work revealed that Dpp4-positive mesenchymal cells contained lysosomal membrane damage and higher SA-β-gal activity. Overall Dpp4 expression in lung was reduced by BPTES treatment in aged mice. Consistent with this, the expressions of Dpp4 and Cadm3 were restricted to C9 (Fig. 1E and 1G).

**Figure 1.**
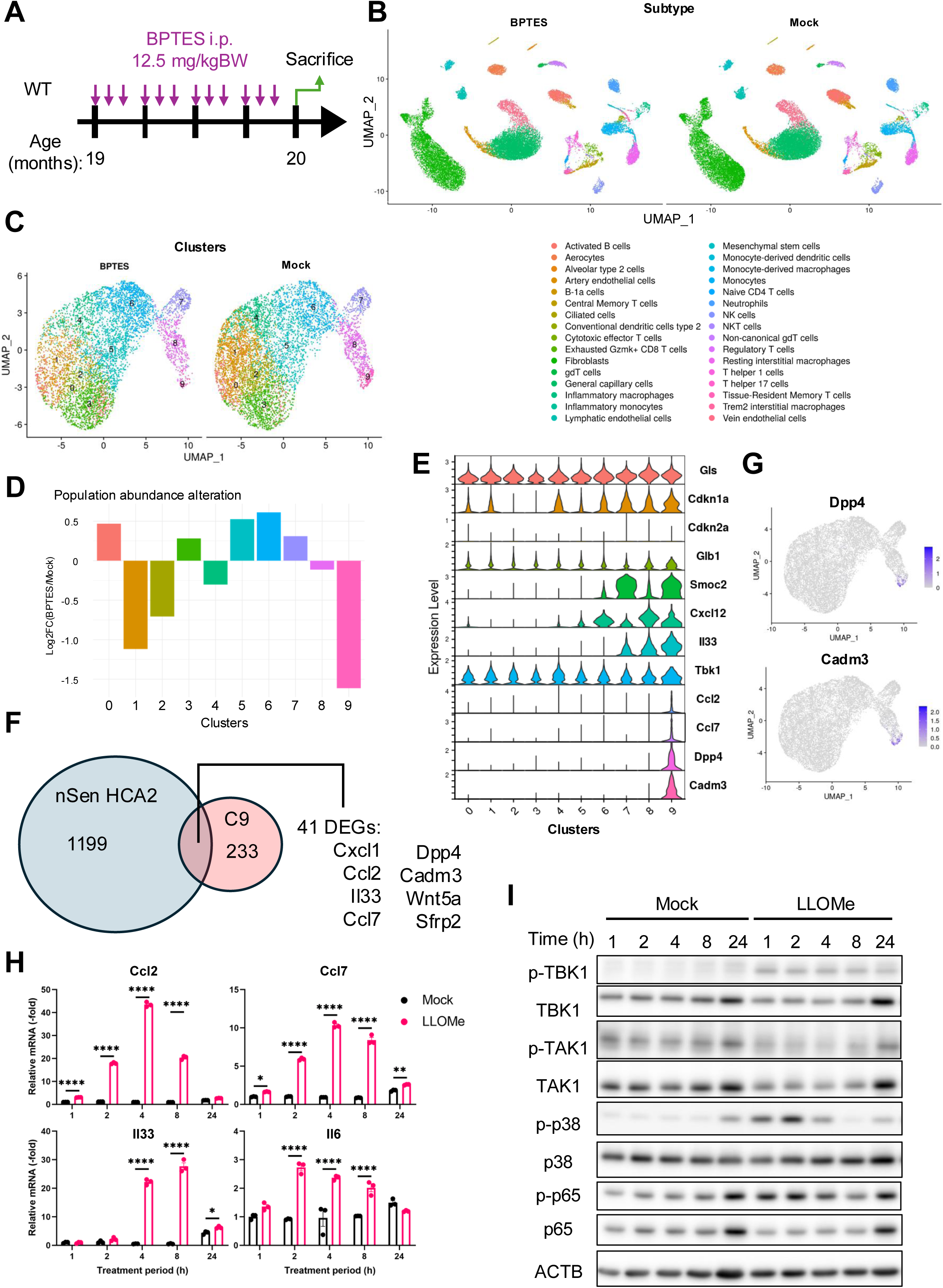
scRNA-seq analysis of lung treated with BPTES revealed the elimination of inflammatory fibroblasts. (A) Schematic diagram showing the schedule of BPTES administration to WT mice from 19-month-old to 20-month-old. (B) UMAP showing the cell types of single-cell transcriptomes obtained from 20-month-old mice treated with BPTES or mock control. (C) UMAP showing the subclusters of pulmonary fibroblasts. (D) Bar plot showing the cell composition alteration in each cluster. Population abundance was calculated as the ratio of the number of cells in each cluster to the total number of cells within the corresponding group. This ratio in the BPTES-treated group was then divided by the ratio in the Mock group, and the log2-transformed value was presented. (E) Violin plot showing the normalized expression levels of the indicated genes in each cluster. (F) The Venn diagram illustrating the overlap between DEGs identified from the re-analysis of RNA-seq data and genes specifically expressed in the C9. Bulk RNA-seq data of nutlin3a-induced senescent (nSen) HCA2 cells were obtained from GSE179465. DEGs in nSen HCA2 cells were defined by Log2FC > 1 and FDR < 0.05. C9-specific genes were identified by comparing the C9 cluster with each of the other clusters individually, selecting genes with Log2FC > 0.3 and FDR < 0.05 in each comparison, and then intersecting the results across all comparisons, resulting in a total of 274 genes. Mouse gene names were converted to human orthologs using MGI Vertebrate Homology. (G) Feature plots showing the normalized expression of indicated genes in pulmonary fibroblasts. (H) qPCR analysis showing the relative expression levels of the indicated genes in primary pulmonary fibroblasts with or without 1 mM LLOMe treatment for the indicated periods. Two-way ANOVA followed by Sidak’s test was performed. (I) Immunoblotting results of pulmonary fibroblasts treated with or without 1 mM LLOMe treatment for the indicated periods by using indicated antibodies.

GSVA revealed that C9 exhibited the highest interferon (IFN) responses and lowest proliferative capacity among all clusters (Fig. S1A). SCENIC analysis suggested the strong transcriptional activities of IRF7, 9 and STAT1 (Fig. S1B). Taken together, these results suggest that the TBK1–IRF signaling pathway was active in C9 [30]; accordingly, Tbk1 expression was highest in C9 among other clusters (Fig. 1E).

### Lysosomal membrane damage induced inflammatory phenotypes in primary cultured pulmonary fibroblasts

We have previously demonstrated that the majority of Dpp4-positive mesenchymal fraction contained lysosomal membrane damage [24]. Thus, we examined whether lysosomal membrane damage could induce inflammatory phenotypes observed in C9. qPCR analysis of LLOMe, a lysosomal membrane damage inducer, -treated mouse primary pulmonary fibroblasts demonstrated the drastic upregulation of Il33, Ccl2, and Ccl7, which were included in overlapped genes shown in Fig. 1E (Fig. 1H). As upstream kinases, the phosphorylation of TBK1 and p38MAPK, but not TAK1, were increased following LLOMe treatment (Fig. 1I). Consistent with this, NF-κ B p65 was also phosphorylated. Therefore, lysosomal membrane damage could activate distinct signaling pathways that trigger the secretion of certain cytokines. This suggests that cells in C9 may harbor lysosomal injury, which could account for their heightened sensitivity to BPTES

### Clusters ablated in p16-DTR model were similar to those targeted by BPTES

p16 is known to be one of the reliable markers for senescent cells. For this reason, various p16-reporter mice were generated and analyzed to characterize senescent cells in vivo. However, there is no clear evidence that these transgenic-based elimination of p16-positive cells target high inflammatory cells. A similar scRNA-seq approach to the BPTES treatment was adopted to address these issues in p16-Cre^ERT2^/Rosa26-lsl-DTR-IRES-tdTomato mice (p16-DTR) [31]. Both aged p16-Cre^ERT2^ and p16-DTR mice (22-month-old) were treated with tamoxifen (TAM) and diphtheria toxin (DT) to selectively remove p16-positive cells in vivo (Fig. 2A). In the lung, similar to the BPTES experiment, we also identified 32 distinct cell types (Fig. 2B). Compared to the p16-Cre^ERT2^ control, we found that the population of cluster 13 (C13) was markedly reduced in p16-DTR with DT treatment (Fig. 2C and 2D). Gene expression pattern of C13 cells shared high similarity with that of C9 in BPTES-treated aged mice (Fig. 1E, 2E, S2A, and S2B). Indeed, the majority (more than 70 %) of differently upregulated genes overlapped between C9 and C13, indicating that C9 in BPTES-treated aged mice and C13 in DT-treated aged p16-DTR mice were corresponding clusters (Fig. 2F). Accordingly, expressions of Dpp4 and Cadm3 (overlapped cell surface markers in 3 groups) were restricted to cluster 13 (Fig. 2G). The C3 was also reduced in p16-DTR group, which was similar to observation of C1 and C2 in BPTES-treated mice. These clusters exhibited the high proliferative potential implying the relatively high plasticity. Thus, eliminating senescent C13 (or C9 in BPTES) clusters might induce the population shift from C3 (or C1 and C2 in BPTES) to other clusters.

**Figure 2.**
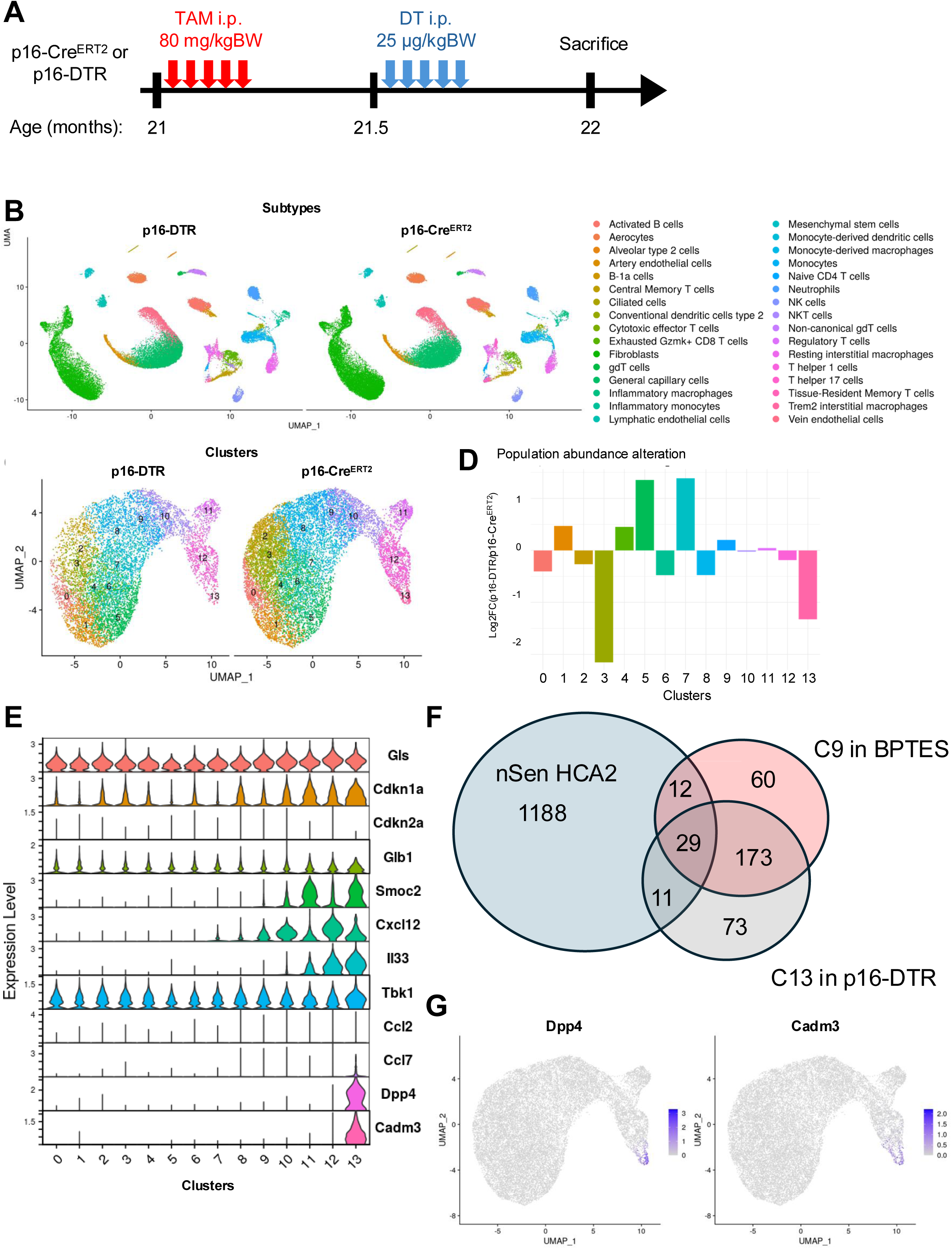
scRNA-seq analysis of lung derived from p16-DTR mice revealed a similar reduction of inflammatory fibroblasts. (A) Schematic diagram showing the schedule of tamoxifen (TAM) and diphtheria toxin (DT) administration to p16-CreERT2 (Control) and p16-DTR mice (elimination of p16-expressing cells) from 21-month-old to 22-month-old. (B) UMAP showing the cell types of single-cell transcriptomes obtained from mice described in A (C) UMAP showing the subclusters of pulmonary fibroblasts. (D) Bar plot showing the cell composition alteration in each cluster. Population abundance was calculated as the ratio of the number of cells in each cluster to the total number of cells within the corresponding group. This ratio in the p16-DTR-treated group was then divided by the ratio in the p16 group, and the log2-transformed value was presented. (E) Violin plot showing the normalized expression levels of the indicated genes in each cluster. (F) The Venn diagram illustrating the overlap between DEGs identified from the re-analysis of RNA-seq data and genes specifically expressed in the C9 (BPTES setting) and C13 (p16-DTR setting). The criteria for gene selection were the same as those described in Figure 1E. (G) Feature plots showing the normalized expression of indicated genes in pulmonary fibroblasts.

### Inflammatory pulmonary endothelial cells were eliminated in both BPTES-treated mice and p16-DTR model

We then analyzed another abundant cell type, pulmonary endothelial cells, with the same approach as that for fibroblasts. Endothelial cells except for aerocytes [32] and lymphatic endothelial cells were subtyped in UMAP (Fig. 3A). Within the general capillary cells, we found the population shift from C4 and C5 to C2 and C3 (Fig. 3B and 3C). Gls expression in C5 was significantly higher than other general capillary clusters (Fig. 3D) (p.adj=1.9e-3). DEG analysis and Gene ontology revealed that cells in C4 and C5 significantly expressed Irf1, Stat1, Stat2, Cx3cr1, Cd24a, and Vav3 and enriched the terms of chemotaxis and cytokine-mediated signaling pathways compared to cells in C2 and C3, implying the chronic inflammatory phenotypes of these clusters (Fig. 3D and 3E) [33–36]. Interestingly, we noticed that C6 showed the highest scores of interferon-related genes in GSVA, which suggests the possible presence of inflammatory cells that are not affected by BPTES (Fig. S3A). We also analyzed the endothelium in p16-DTR approach (Fig. 3F). The population shifted from C4 to C1 and C2 following the elimination of p16-positive cells, whose direction was different to the patterns in BPTES treatment (Figs. 3B, 3G, and 3H). By comparing C4 with C1 and C2, we observed that C4 enriched cytokine-related terms and additionally showed elevated expression of angiogenesis-related genes. Similar to the BPTES experiment, the p16-DTR experimental design also revealed a population of cells (C5) that remained unaffected by DT treatment and expressed the highest levels of interferon-related genes (Fig. S3B). Both approaches suggest the existence of a subset of general capillary cells that lack lysosomal membrane damage and does not express p16, contributes to the inflammatory phenotype.

**Figure 3.**
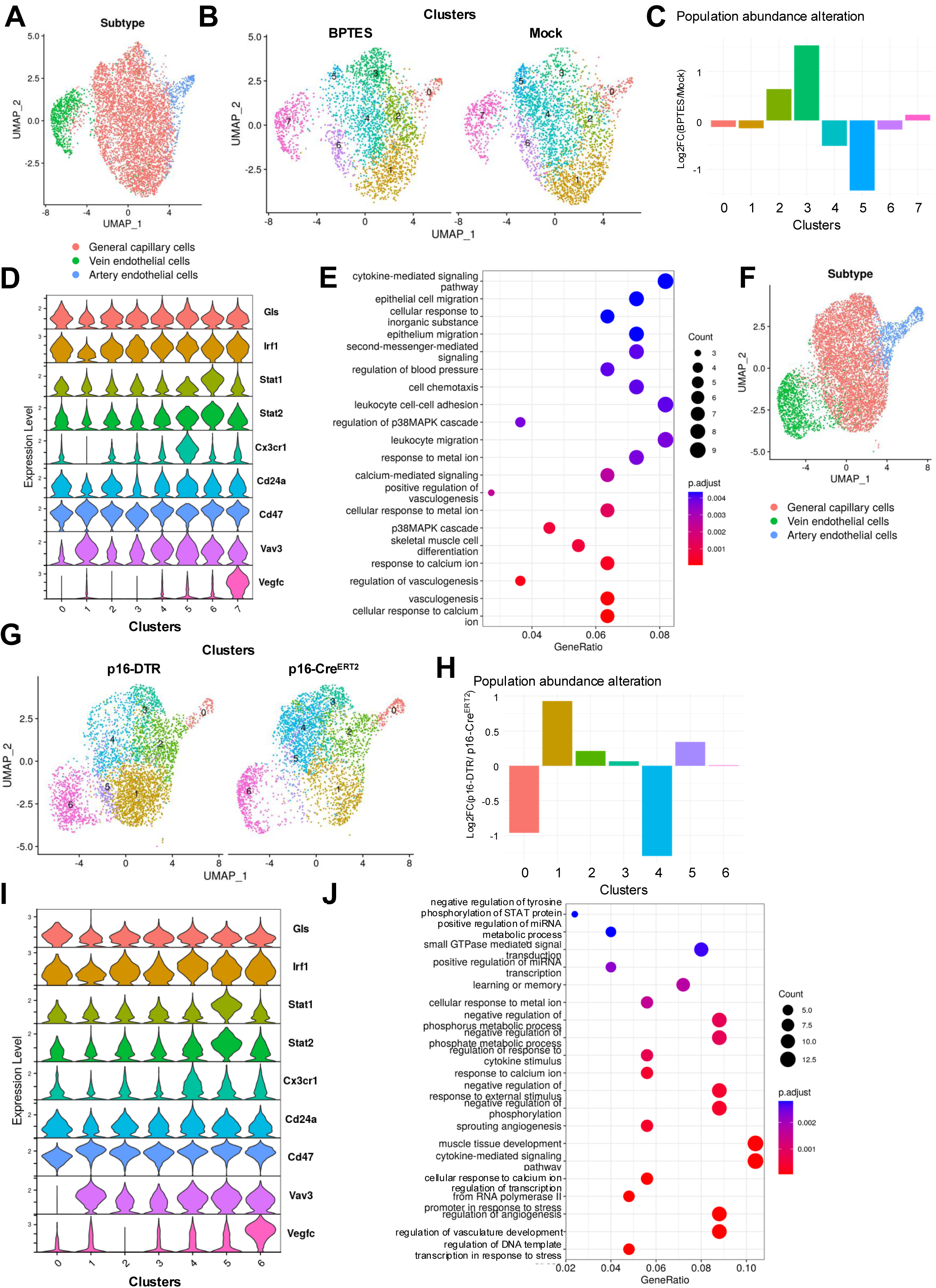
scRNA-seq analysis of pulmonary vascular cells exhibited the partial amelioration in inflammatory phenotypes by BPTES or DT treatments. (A, B) UMAP showing the (A) subtypes and (B) subclusters of endothelial cells in BPTES setting. (C) Bar plot showing the cell composition alteration in each cluster. (D) Violin plot showing the normalized expression levels of the indicated genes in each cluster. (E) Dot plot showing the significantly enriched GO terms in up-regulated DEGs identified by comparing C4 and C5 with C2 and C3. All DEGs were determined by log2FC > 0.3 and FDR < 0.05, and the threshold of GO terms was also FDR < 0.05. (F, G) UMAP showing the (F) subtypes and (G) subclusters of endothelial cells in p16-DTR setting. (H) Bar plot showing the cell composition alteration in each cluster. (I) Violin plot showing the normalized expression levels of the indicated genes in each cluster. (J) Dot plot showing the significantly enriched GO terms in up-regulated DEGs identified by comparing C4 with C1 and C2. All DEGs were determined by log2FC > 0.3 and FDR < 0.05, and the threshold of GO terms was also FDR < 0.05.

### BPTES treatment altered immune cell properties in lungs

In addition to non-immune cells, immune cells are also involved in chronic inflammation. We separated the immune cells into myeloid-lineage and T-lineage population, and then annotated the subtypes based on the typical markers. In the T-lineage cells, the population of cytotoxic effector T (CTL), natural killer cells (NK), and central memory T (TCM) cells were reduced in BPTES-treated group. The former two cell types are known major donors of IFNs, which contribute to inflammaging (Fig. 4A and 4B) [37]. Although we found the reduction in these certain cell types, it should be noted that the expression of Gls were fairly consistent across T subtypes (Fig. 4C).

**Figure 4.**
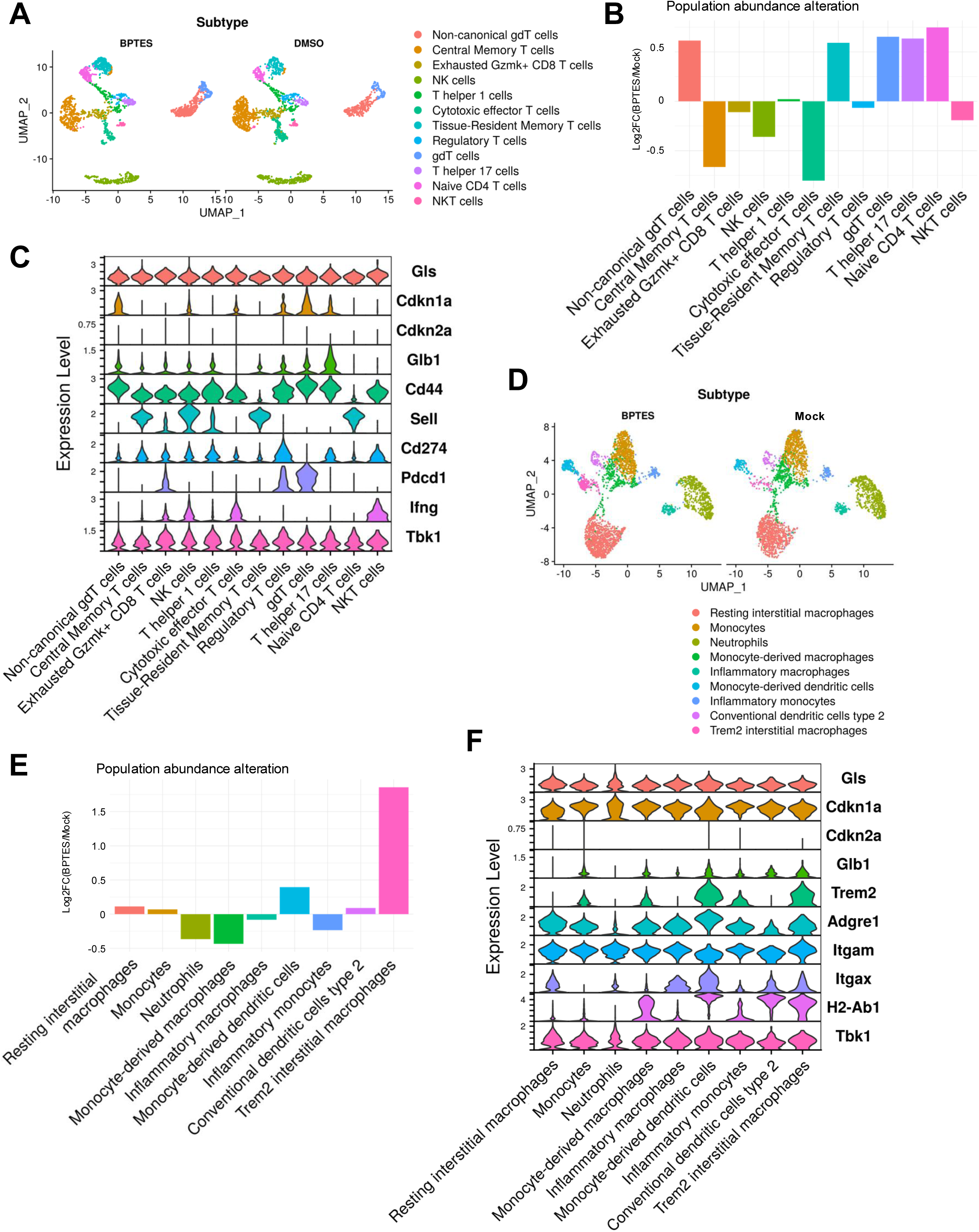
Immunoactivities of immune cells were repressed in mice treated with BPTES. (A) UMAP showing the subtypes of T-lineage cells in BPTES setting. (B) Bar plot showing the cell composition alteration in each subtype. (C) Violin plot showing the normalized expression levels of the indicated genes in each subtype. (D) UMAP showing the subtypes of myelocyte-lineage cells in BPTES setting. (E) Bar plot showing the cell composition alteration in each subtype. (F) Violin plot showing the normalized expression levels of the indicated genes in each subtype.

In myeloid-lineage cells, the population of Trem2-positive macrophages were drastically increased in BPTES-treated group (Fig. 4D and 4E). Trem2-positive macrophages have been reported to suppress inflammation induced by other immune cells [38–40]. Again, we found that the expression of Gls were fairly consistent across macrophage subtypes (Fig. 4F). Therefore, we assume that the impact of BPTES on the immune microenvironment was unlikely to result from direct cytotoxic effects on immune cells. Rather, it is more likely that the senolytic activity of BPTES toward inflammatory non-immune cells indirectly modulates the inflammatory strength of immune cells.

To evaluate the cross-communication between inflammatory non-immune cells and other immune cells, we reassigned the fibroblast in the Mock sample into 5 groups and named the cells in C9 as Pi16_Sene (Fig. S4A). We then performed cell-cell communication analysis using CellChat to calculate the ligand-receptor pair interactions among all subtypes within the Mock sample [41]. All the ligands secreted by fibroblasts were included in our analysis pipeline, and we selected the ligands with the altered expression in these 5 groups of fibroblasts (Fig. S4B). Based on the results, we found that Nampt, Wnt11, Ccl11, and Ccl19 were prominently expressed in both Pi16_Sene and the neighboring Pi16_B clusters. In addition, Ccl2, Ccl7, Il18, Plau, Sema3c, and Lgals9 were specifically expressed in Pi16_Sene cluster and were inferred to regulate various other cell types, including endothelial cells, myeloid-lineage cells, and T lineage cells (Fig. 5). Among them, CCL19, IL18, and CCL7 have been demonstrated to be involved in T cell recruitment, pathogenic pneumonia, and pulmonary fibrosis induced by systemic sclerosis, respectively [42–44]. As mentioned above, among pulmonary fibroblasts, the Pi16_Sene population interacted with other cell types more extensively through cytokine secretion than did the other clusters; these signaling pathways encompassed many ligands related to pulmonary inflammation and fibrosis. Moreover, this cell population was markedly diminished following BPTES treatment, which aligned with the observed decrease in pro-inflammatory T cells and increase in immunosuppressive Trem2-positive macrophages in the lungs of BPTES-treated mice.

**Figure 5.**
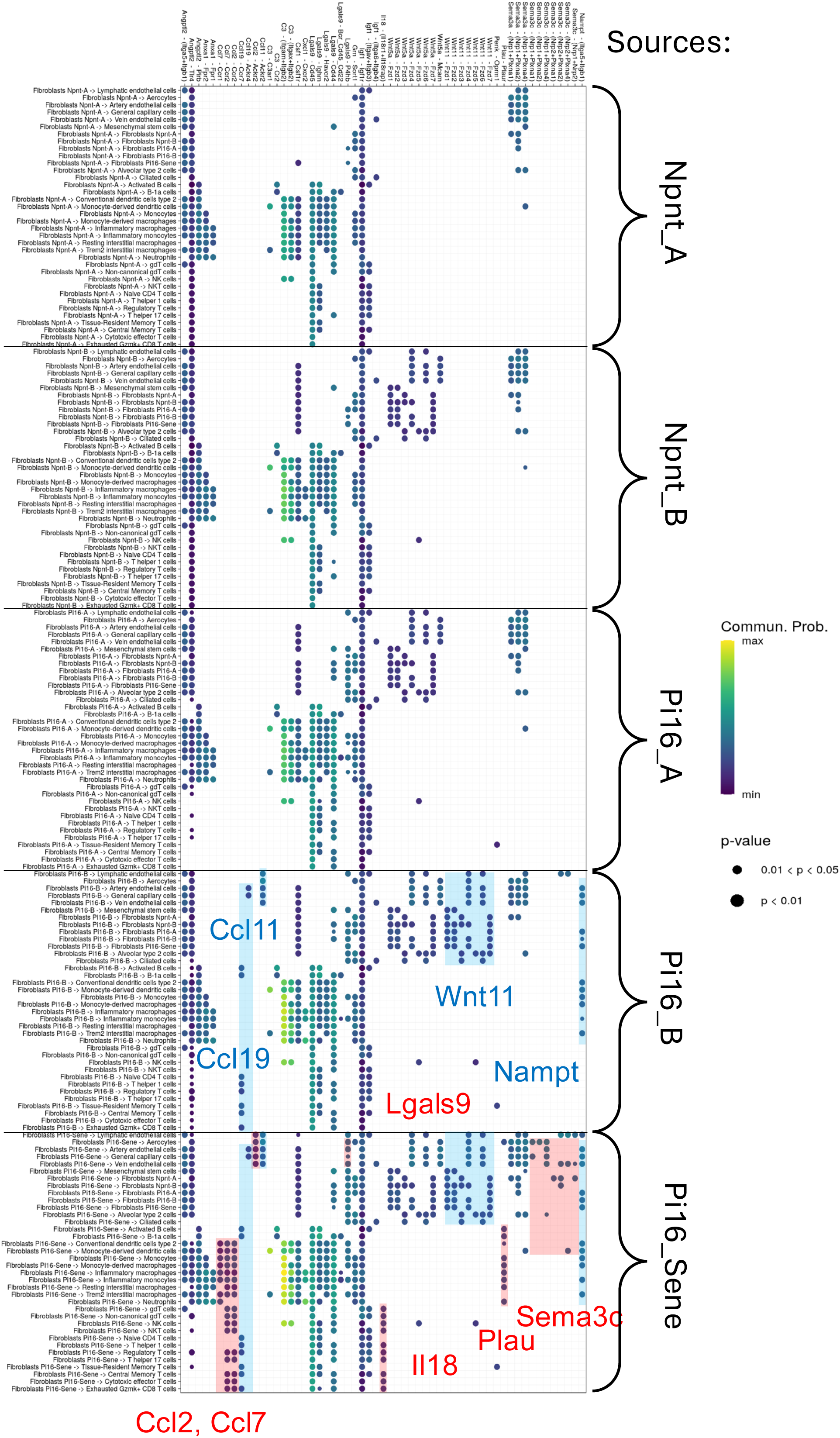
Cell-cell communication inference showed the strong impact of inflammatory fibroblast cluster which sensitized to BPTES. The dot plot displays all significantly enriched ligand-receptor pairs from fibroblasts to other subtypes. The color of each dot represents the communication probability. Blue rectangles highlight ligand-receptor pairs specific to both Pi16_B and Pi16_Sene, while red rectangles indicate those unique to Pi16_Sene. The fibroblast subpopulations used for CellChat analysis were annotated and displayed in Figure S4A, and ligands highly expressed in Pi16_Sene were incorporated into the analytical pipeline.

### BPTES treatment eliminated inflammatory injured proximal tubular cells in aged kidney

To examine the BPTES effects in other organs, we performed the scRNA-seq analysis on the kidney isolated from the same mice. A total of 34 cell types were identified by marker genes and visualized in the UMAP (Fig. 6A). Proximal tubule (PT) mainly maintains the balance of fluid, electrolytes, and nutrients, and has been widely reported to involve in the progression of renal pathology [45]. The PT cells could be further characterized as proximal convoluted tubule (PCT), proximal straight tubule (PST), and injured proximal tubule (injured-PT) by the expressions of Slc5a2, Atp11a, and Dcdc2a, respectively (Fig. 6B and S5A) [46]. Among the PCT, BPTES treatment led to the population shift from C0 to C3, and the drastic reduction in the population of injured-PT (Fig. 6C and 6D). Injured-PT expressed the high levels of Gls (p.adj=5.14e-15), Nfkb1, Stat1, and Irf9, which revealed the inflammatory phenotype and explained the eliminating effect by BPTES. Besides, cells in C0 mildly but expressed higher levels of Gls (p.adj=1.75e-6) and IFN response-related genes (p.adj=5.08e-29) than cells in C2 and C3 (Fig. 6E and S5B). Based on GSVA and SCENIC, we found that injured-PT showed the highest scores of inflammatory-related terms and -dominant transcription factors (Fig. S5B and S5C). On the other hand, PCT cells in C0 exhibited transcriptional activities of IRFs compared to cells in C3, which suggests the improvement of local chronic inflammation in the microenvironment of tubular structure (Fig. S5C) (IRF9 p.adj=5.85e-24). Consistent with this, the GSVA scores of IFN responses and other key inflammatory pathways showed the overall decrease in BPTES group rather than Mock group (Fig. 6F).

**Figure 6.**
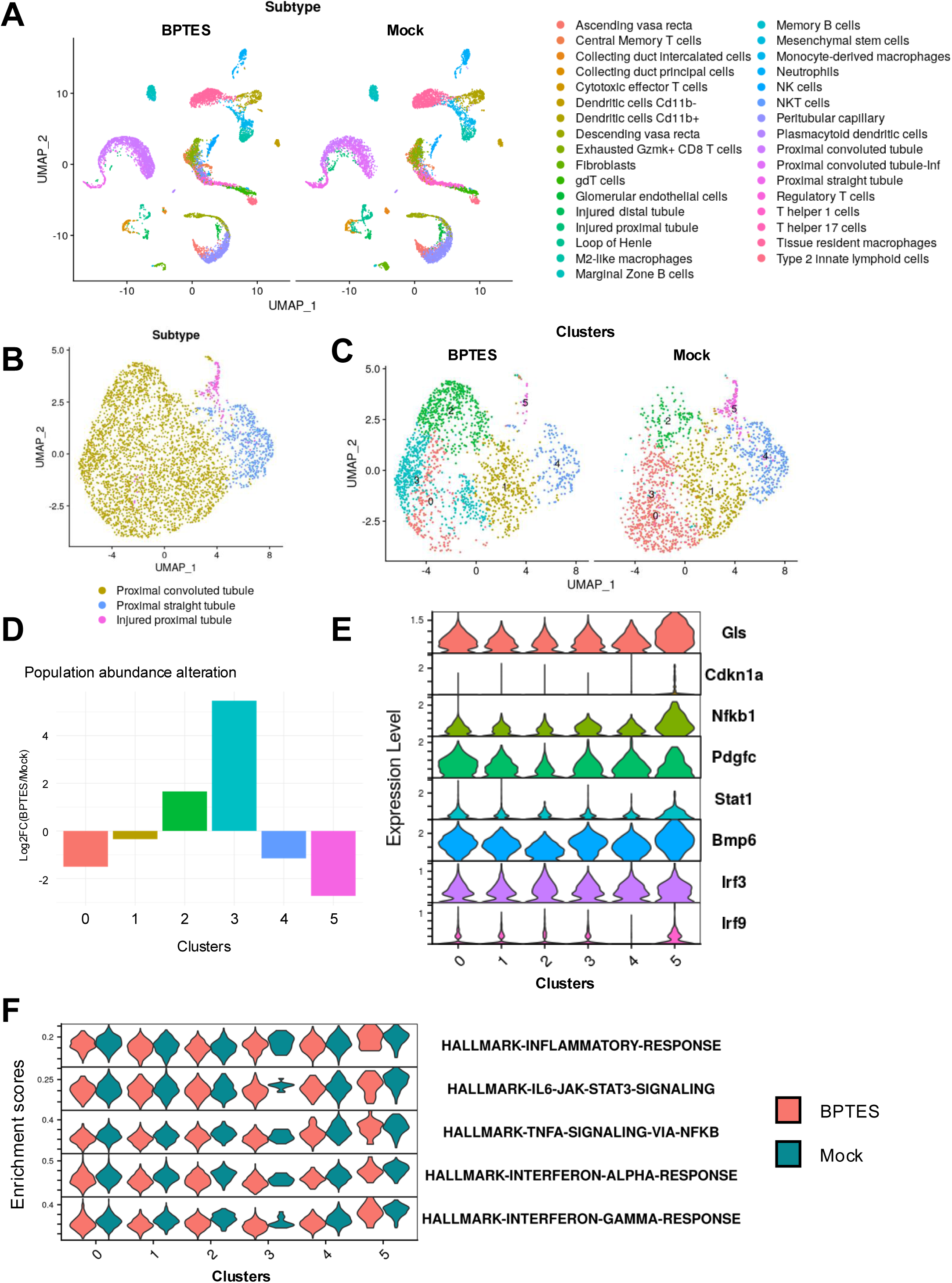
scRNA-seq analysis of kidney treated with BPTES revealed the elimination of injured and inflammatory proximal tubular cells. (A) UMAP showing the cell types of single-cell transcriptomes obtained from 20-month-old mice treated with BPTES or mock control. (B, C) UMAP showing the (B) subtypes and (C) subclusters of proximal tubular cells. (D) Bar plot showing the cell composition alteration in each cluster. (E) Violin plot showing the normalized expression levels of the indicated genes in each cluster. (F) Violin plots showing the enrichment scores of indicated terms analyzed by GSVA.

### Injured-PT was the hub of inflammatory signaling network in aged kidney

To further investigate the cell-cell communication between PT and immune cells, we conducted the Cellchat analysis on transcriptomes in Mock group [41]. Because PCT cells in C0 exhibited higher inflammatory phenotypes than other clusters of PCT cells, which were reassigned as PCT_inf followed by further analysis. According to the analysis results, we noticed that several growth factors such as Egf and Pdgfb were secreted by PCT_Inf, which were related to renal fibrosis and injury [47–49]. In addition, Injured-PT cells secreted Il34, Cxcl16, Clcf1, and Sema3c, which activated macrophages, T cells, renal fibrosis and correlated with kidney injury [50–53] (Fig. S6). These results suggest that the PT cells eliminated by BPTES treatment were the major contributors to the pathogenesis of age-dependent kidney dysfunction, which highly imply the ameliorative effects of GLS1 inhibitor.

## Discussion

The benefits of senolytic treatment have been widely accepted to improve certain age-related dysfunction and diseases. However, the exact targeting populations in vivo and their characteristics remain largely unknown. This point is a critical barrier to the development of senolytic therapies for age-related diseases. In this study, we obtain robust evidence that GLS1 inhibitor, BPTES, eliminated the highly inflammatory non-immune cells in both lung and kidney by unbiased single-cell transcriptome approach. In particular, pulmonary fibroblasts in C9 eliminated by BPTES shared the general characteristics with in vitro senescent culture fibroblasts. There are several pieces of evidence: (1) High transcriptional activity of IRF family and inflammatory signatures. (2) Dpp4, a known senescent fibroblast marker [24,29,54], expression was restricted to the C9 cluster. (3) Intersection of up-regulated DEGs between human senescent foreskin primary fibroblasts, HCA2, and fibroblasts in C9 mainly composed of inflammation-related genes. (4) Cells in C9 expressed the highest levels of Glb1 encoding SA-β-gal, whose strong activity was previously demonstrated by staining on sorted DPP4-positive fibroblasts. (5) Score of proliferative potential-related terms such as “mitotic spindle” was the lowest in C9. (6) Fibroblasts eliminated by DT in p16-DTR mice were corresponding to cells in C9, confirmed by transcriptomic profiles. These results clearly proved the senolytic effect of BPTES in fibroblast population.

It should be noted that DPP4-positive mesenchymal cells had been shown to predominantly suffer from lysosomal membrane damage, at least, in the lung, skin, and white adipose tissues [24]. Lysosomal membrane damages as well as Dpp4 expression were possibly the common features of senescent fibroblasts in vivo independent of organs presented. In addition, we also found other types of non-immune cells, such as pulmonary general capillary cells and proximal tubular cells showing inflammatory properties, which were also sensitive to BPTES.

As for pulmonary endothelial cells, a part of inflammatory cells was also eliminated by BPTES (C5) and DT (C4). These results suggest that lysosomal membrane damages were correlated with inflammatory phenotypes even in the endothelium. It has been reported recently that lysosomal dysfunction was observed in the arterial endothelium of pulmonary arterial hypertension (PAH) [55]. In fact, the senescent pulmonary fibroblasts identified in this study were Pi16-positive, which has been reported to localize in the adventitial cuff and contribute to PAH by modulating the immune cell infiltration [56–58]. Thus, lysosomal membrane damages in the pulmonary stromal cells might play a key role in detrimental lung diseases. Interestingly, we identified another subpopulation (C6 in BPTES setting) showing the high IFN signatures resistant against BPTES as well as DT (p16-DTR mice) treatments. So far, the impact of this population on age-related inflammation has still been unexplored.

Innate and adaptive immunity are also indispensable for establishing chronic inflammatory microenvironment [59]. In this context, we revealed that cells eliminated by BPTES made a significant contribution to the overall inflammatory signaling networks. In the lung, several cytokines expressed by Pi16_Sene fibroblasts were significantly enriched to stimulate macrophages and T lymphocytes. Loss of these signals may explain the molecular basis underlying immune cell composition alteration from cytotoxic T cells to relatively resting status. Besides, the expansion of Trem2-positive macrophages also reflected the immunosuppressive niche, although their long-term presence may induce fibrosis [39]. The influence of these fibroblasts involves various cytokines, and thus blocking the single signal may be insufficient for suppressing the chronic inflammation. In the kidney, injured-PT cells exhibited distinctively strong inflammatory signatures and Gls expression, which was sensitive to BPTES. This population interacted with immune cells and fibroblasts through secreting multiple pro-inflammatory and fibrogenic cytokines, known to be related to kidney diseases.

With respect to the clinical application of GLS1 inhibitors to age-related diseases, it has been reported that BPTES treatment can ameliorate age-related phenotypes in kidneys such as renal fibrosis [25]. Another GLS1 inhibitor, CB-839, is shown to improve PAH symptoms [27] and pulmonary fibrosis [26]. Although these previous reports did not examine in depth the effects of GLS1 inhibition on inflammation, given that perivascular lysosomal dysfunction is indeed increased in PAH models [55], both pieces of evidence clearly highlight the therapeutic potential of GLS1 inhibitors.

## Methods

### Mouse experiment

Mice were housed in a temperature (23-25°C), and humidity-controlled colony room, maintained on a 12-h light/dark cycle (08:00 to 20:00 light on), with standard food (CA-1, CLEA Japan), and water provided ad libitum. All animals were handled following the Guidelines for Animal Experiments of the Institute of Medical Science, the University of Tokyo, and the Institutional Laboratory Animal Care. p16-DTR mice were generated by crossing p16Ink4a-CreERT2 mice [31] with Rosa26-SA-lsl-DTR-IRES-tdTomato mice [60]. All p16-DTR mice and littermate control, p16-CreERT2 mice, were daily i.p. injected with tamoxifen (TAM, 80 mg/ kgBW) and diphtheria toxin (DT, 25 μg/ kgBW) for five days at the designated time point. BPTES dissolved in DMSO were diluted 10 times in corn oil and i.p. injected into C57BL/6 mice with 0.25 mg/ 20 gBW/ 200 ul. 19-month-old mice were administered with BPTES or Vehicle control 3 times per week over a 4-week period.

### Culture of primary pulmonary fibroblasts

Lung tissues were harvested from anesthetized wild-type mice and finely minced using surgical blades. The minced tissue was then incubated to enzymatic digestion in a solution containing Liberase TM (50Uμg/mL, Roche), HEPES (10UmM, nacalai tesque), and DNase I (2 kunitz/mL, Sigma) in RPMI 1640 medium (nacalai tesque). The digestion was placed in 37°C and carried out in three 20-minute intervals, during which the tissue was mechanically disrupted using repeated pipetting with 18G and 21G needles. After digestion, the cell suspension was filtered through a 70Uμm cell strainer following the addition of RPMI containing 10% FBS, and red blood cells were lysed using RBC Lysis buffer (Thermo Fisher Scientific). The resulting cells were cultured in a 5% O₂ hypoxia incubator for three days. Cells were then detached with trypsin, washed with PBS, and sorted by FACS to isolate DAPI⁻/CD31⁻/CD45⁻/CD140a⁺ cells for further culture and experiments. LLOMe was applied at a concentration of 1UmM for the indicated period.

### Immunoblotting

Whole-cell lysates were prepared by directly lysing cells in Laemmli buffer (2% SDS, 10% glycerol, 5% 2-mercaptoethanol, 0.002% bromophenol blue, and 62.5 mM Tris-HCl, pH 6.8). Proteins were separated by SDS-PAGE, transferred to a PVDF membrane (Immobilon-P; Millipore), and subjected to immunoblotting using the indicated antibodies and an ECL detection system. An ACTB antibody (sc-47778, Santa Cruz) was used as a loading control. Images were acquired with an Amersham Imager 680 (Cytiva). All antibodies used in this study contained: p65 (CST 8242), phospho-p65 Ser536 (CST 3033), TAK1 (CST 5206), phospho-TAK1 Ser412 (CST 9339), TBK1 (CST 3504), phospho-TBK1 Ser172 (CST 5483), p38 (CST 9212), and phospho-p38 Thr180/Tyr182 (CST 9216).

### RNA extraction and quantitative PCR

RNA extraction was performed according to the manufacturer’s protocol using the RNeasy Mini Kit (QIAGEN). For cDNA preparation, reverse transcription was carried out using the ReverTra Ace qPCR RT Master Mix (TOYOBO), following the manufacturer’s protocol. Gene expression was measured using quantitative real-time PCR (qPCR) using cDNA as a template. qPCR reactions were conducted using THUNDERBIRD SYBR qPCR Mix (TOYOBO) with ROX reference dye in 96-well optical reaction plates. The relative expression of each gene was calculated by the Delta-Delta Ct method using Actb for normalization. All primers are listed in Table S1.

### scRNA-seq library preparation

The mice were anesthetized and further systemically perfused with PBS to remove the circulating cells. Lungs and kidneys were collected, minced, and incubated for 20 minutes in 50 μg/mL Liberase TM (for lung) or TH (for kidney), 10 mM HEPES, 2 kunitz/mL DNase in RPMI at 37°C and passed through 18G needles followed by 21G needles with a 20 min incubation in between. Cells were filtered through a 70 μm cell strainer and resuspended into RBC lysis buffer for 3 min at room temperature and washed with wash buffer. Cells were treated with an FcR blocking reagent (Miltenyi Biotec) on ice for 10 min, stained with anti-CD45 at 4°C for 30 min, washed, and sorted using FACS AriaIII (BD Biosciences).

### scRNA-seq analysis

For the scRNA-seq library construction, we followed the manufacturer’s instructions of 10x Genomics Chromium GEM-X Single cell 3′ Reagent Kit v4. The single-cell suspensions of DAPI-/CD45+ and DAPI-/CD45-cells were sorted from the tissue lysates with 9,000 cells and 20,000 cells, respectively. Library sequencing was performed on the DNBSEQ-G400RS (MGI Tech) with 150 bp paired-end reads. The Cell Ranger package (version 8.0.1) was utilized to process unique molecular identifiers (UMIs) and barcodes and align the transcripts to a mm10 mouse reference genome. After obtaining the feature-barcode matrix, we utilized Seurat package (ver. 4.2.0) in R language (ver. 4.2.1) to process the quality control, clustering, dimension reduction, cell type annotation, and identification of differentially expressed genes (DEGs). The thresholds of quality control included 2000<nFeatures<7500, nCounts<50000, and mitochondrial counts ratio<5% (for lung) or <20% (for kidney). After quality control filtering, all single-cell transcriptomes were selected for further normalization, log transformation, dimensional reduction, and clustering. Marker genes were generated by the FindAllMarkers function using the Wilcoxon rank-sum test, and then cell clusters were assigned to specific cell populations based on the expression of canonical markers of these cell populations.

The DEGs between each cluster were identified by using the FindMarkers function. The two-sided Wilcoxon rank-sum test followed by B-H method was conducted to calculate the adjusted p-values for each identified DEG with Log2FC > 0.3 and adjusted p-values < 0.05. Gene Ontology (GO) analysis to identify the enriched terms of biological processes by packages clusterProfiler (ver. 4.6.2) [61]. The significance of all terms was identified by adjusted p-value<0.05 using B-H method.

The gene set variation analysis (GSVA) was performed to estimate the enrichment of annotated mouse gene sets obtained from MSigDB for each cell using GSVA (ver. 1.47.3) [62]. The transcriptional regulatory inference of scRNA-seq datasets was examined using the SCENIC (version 1.3.1) workflow [63], specifically employing the GENIE3 (version 1.20.0) [64] and RcisTarget (version 1.18.2) R packages, with default parameters. The reference transcription factors (TFs) for the mm10 genome were obtained using RcisTarget.

The cell-cell communication inference was performed by using CellChat (version 2.2.0) package [41]. The cell type annotated Seurat object was first transformed into the CellChat object and then followed the default analysis pipeline to process the ligand-receptor determination. The “Secreted Signaling” reference ligand-receptor datasets were conducted in the inference.

## Data availability

All materials are available from corresponding authors on request. The scRNA-seq datasets described in this article have been uploaded to Gene Expression Omnibus (GEO) with accession number GSE300530, GSE300532, and GSE301292.

## Acknowledgments

We are grateful to Mrs. Chieko Konishi, Yoshie Chiba, and Tomoko Ando for their technical assistance. The super-computing resource was provided by the Human Genome Center (University of Tokyo). This study was supported by Pathology Core Laboratory and FACS Core Laboratory, Institute of Medical Science, University of Tokyo. Computational resources were provided by the supercomputer system SHIROKANE at the Human Genome Center (Univ. of Tokyo). This study was supported by AMED under Grant Numbers 21zf0127003 (M.N.), 21cm0106175 (M.N.), and 21gm5010001 (M.N.), 214600040 (Y.J.), and by MEXT/JSPS KAKENHI under Grant Numbers 20H00514(M.N.), 19H05740 (M.N.), JP18H05026m (Y.J.), JP16H06148 (Y.J.), JP16K15238 (Y.J.), 23KJ0711 (Y.Y.O.), 25K18870 (T-W.W.) and by the Princess Takamatsu Cancer Research Fund (M.N.).

---

**Figure S1.**
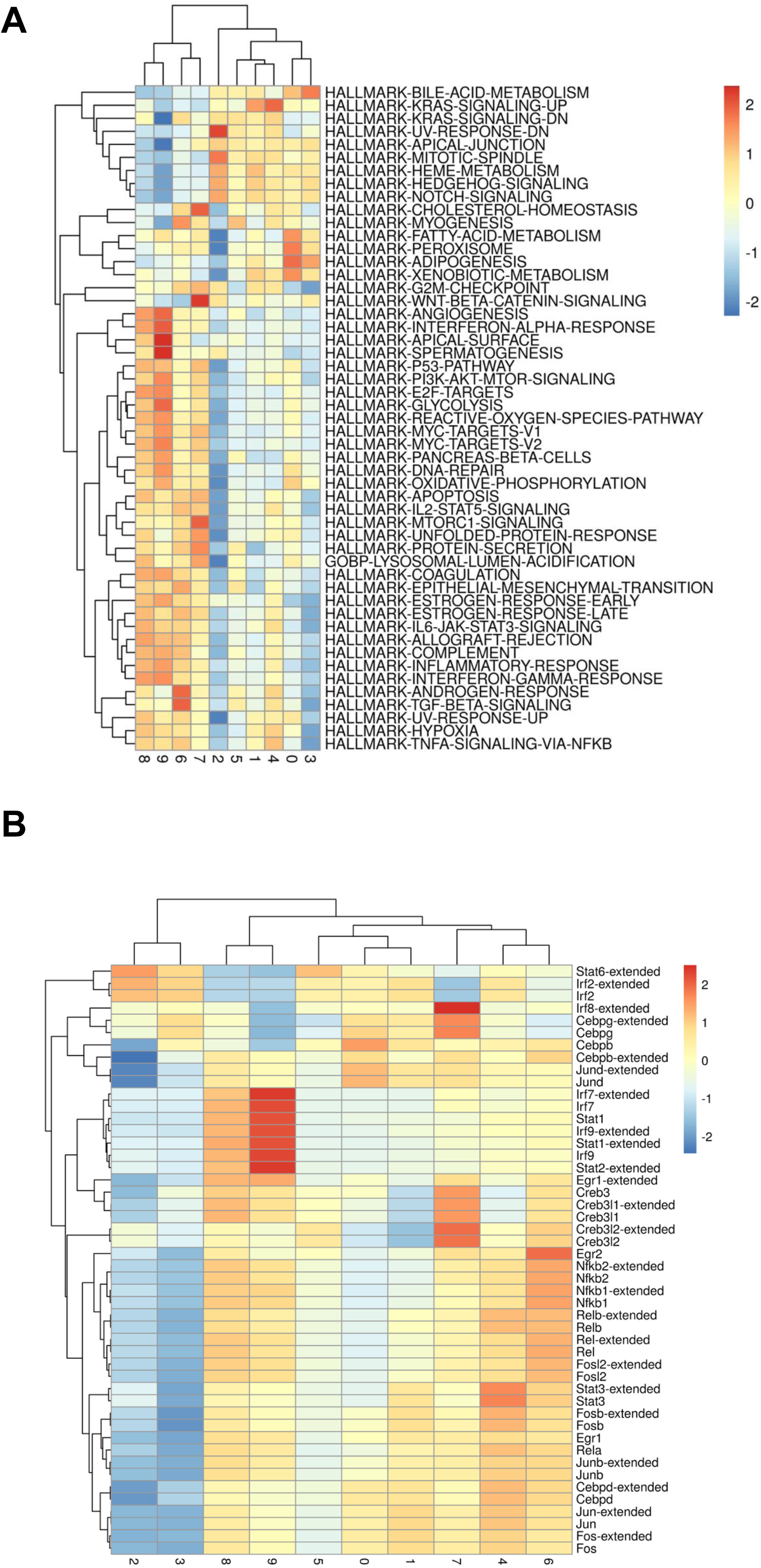
scRNA-seq analysis of lungs treated with BPTES revealed the elimination of inflammatory fibroblasts. (A, B) Heatmap showing the average (A) GSVA scores and (B) SCENIC scores of indicated terms or transcription factors in each cluster of pulmonary fibroblasts from BPTES scRNA-seq datasets.

**Figure S2.**
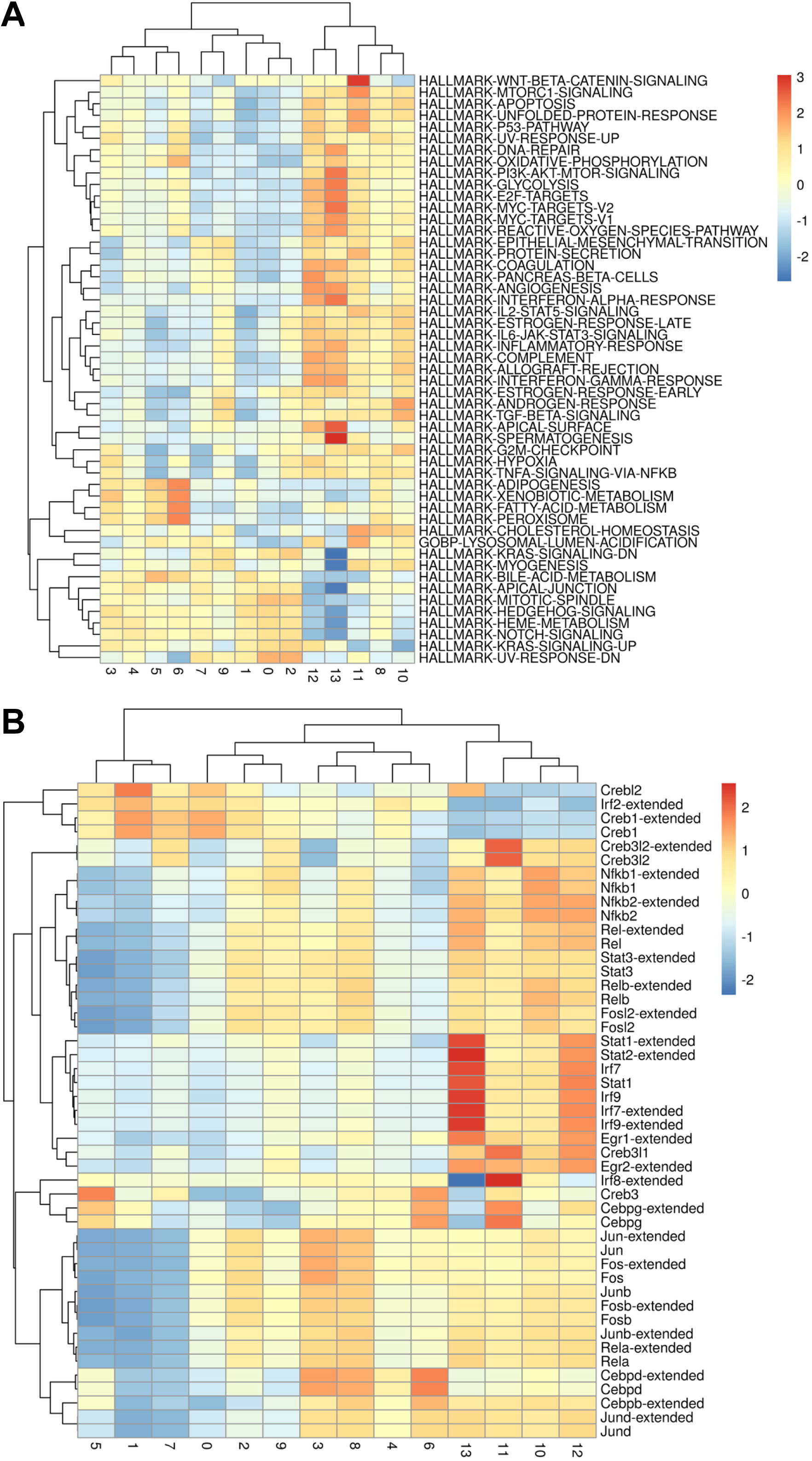
scRNA-seq analysis of lung derived from p16-DTR mice revealed a similar reduction of inflammatory fibroblasts. (A, B) Heatmap showing the average (A) GSVA scores and (B) SCENIC scores of indicated terms or transcription factors in each cluster of pulmonary fibroblasts from p16-DTR scRNA-seq datasets.

**Figure S3.**
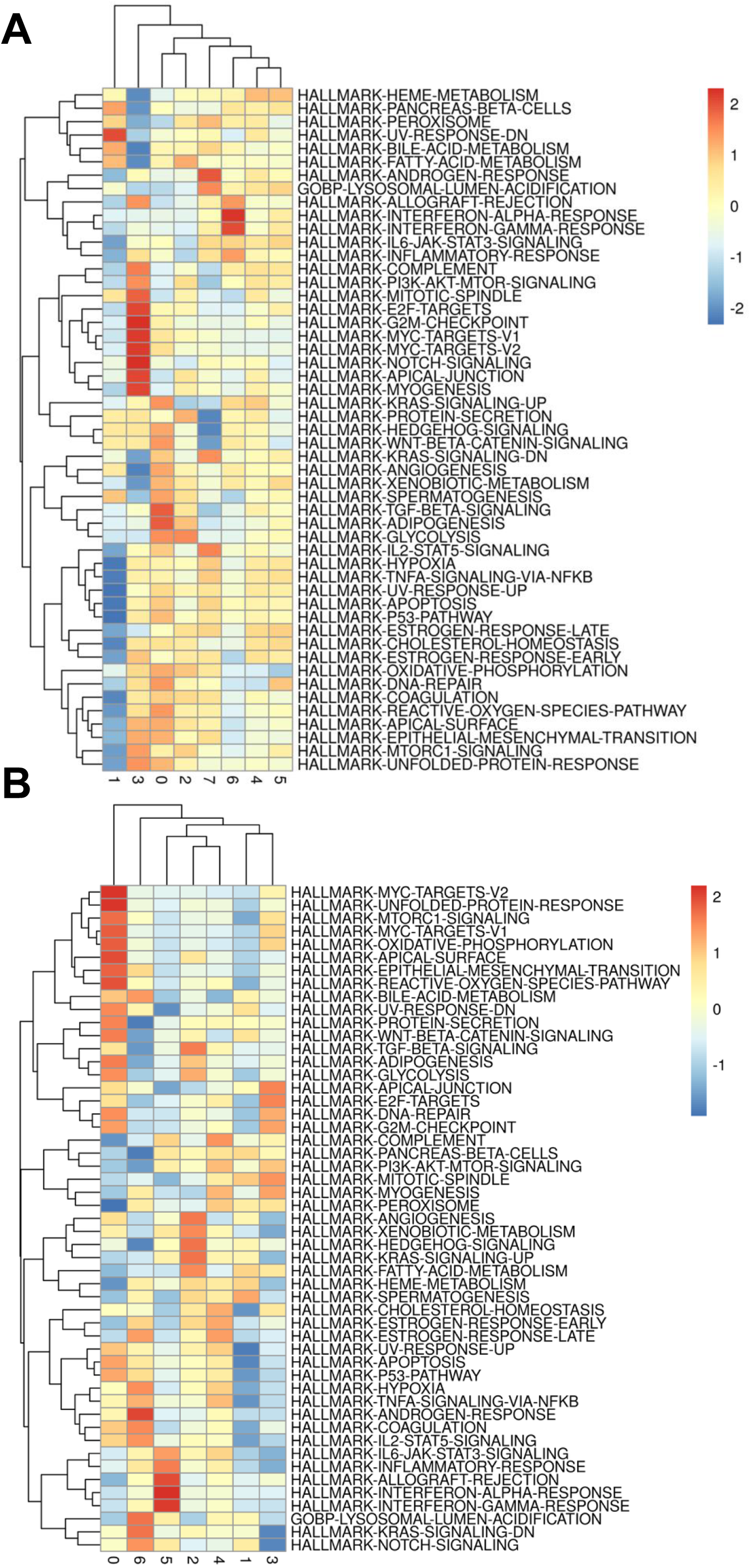
scRNA-seq analysis of pulmonary vascular cells exhibited the partial amelioration in inflammatory phenotypes by BPTES or DT treatments. (A, B) Heatmap showing the average GSVA scores of indicated terms in each cluster of pulmonary vascular cells isolated from (A) BPTES and (B) p16-DTR scRNA-seq datasets.

**Figure S4.**
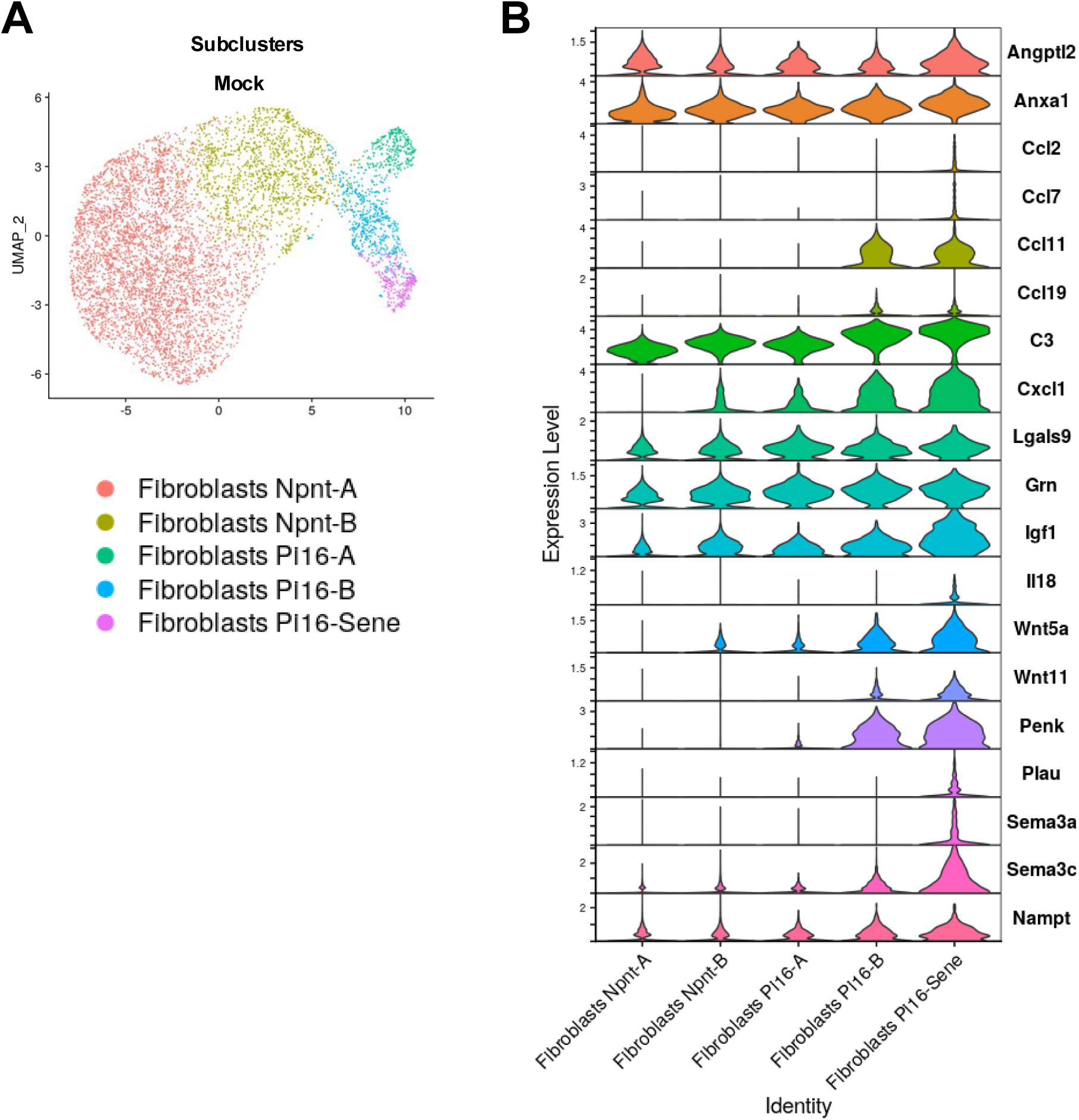
Inflammatory fibroblast cluster expressed higher levels of several cytokines. (A) UMAP showing the subclusters of pulmonary fibroblasts from Mock-treated mice used in Cell chat analysis. (B) Violin plots showing the normalized expression levels of indicated genes in each cluster described in A

**Figure S5.**
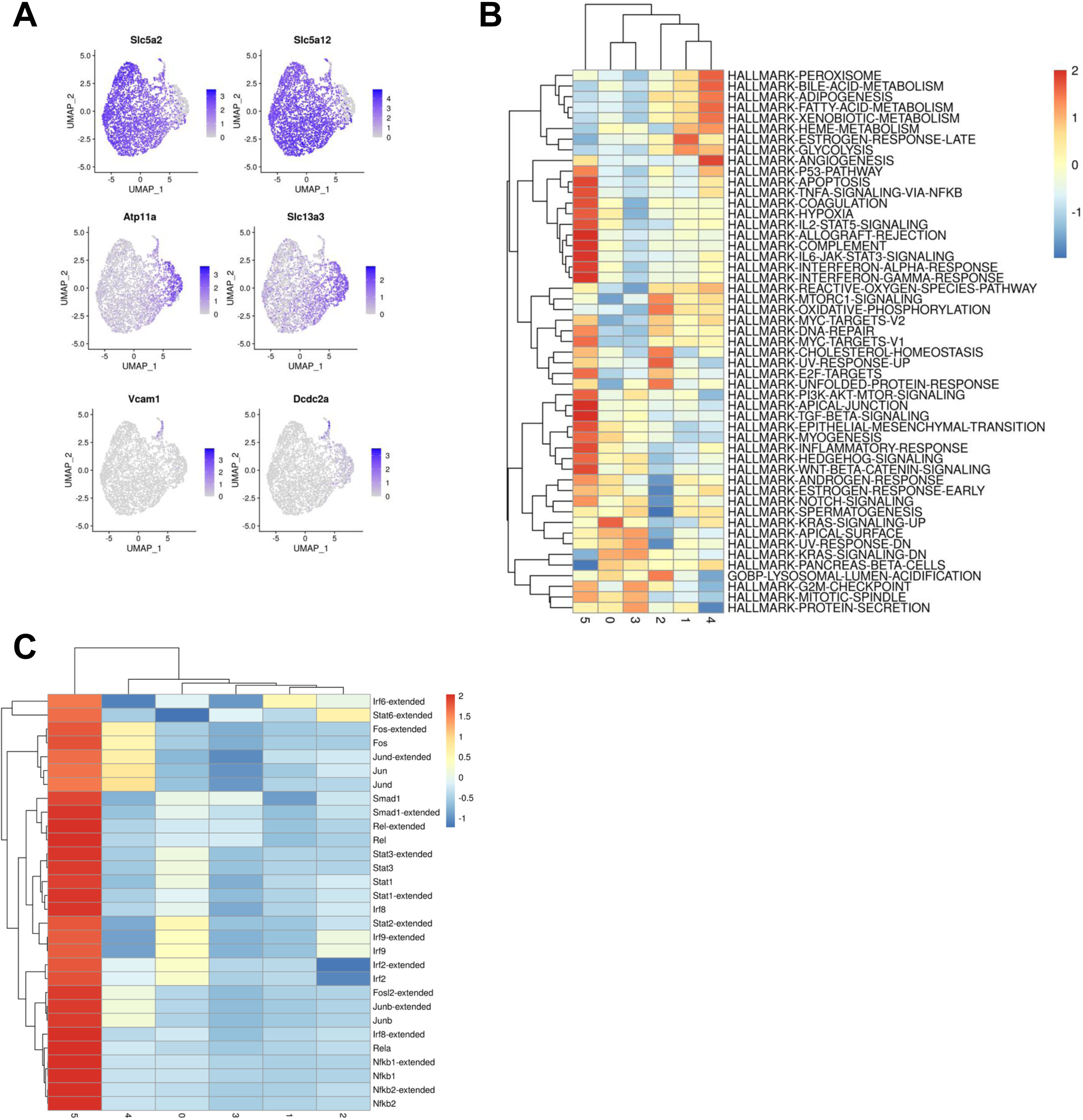
scRNA-seq analysis of kidney treated with BPTES revealed the elimination of injured and inflammatory proximal tubular cells. (A) Feature plots showing the expression levels of marker genes used in distinguished subtypes of proximal tubular cells. (B, C) Heatmap showing the average (B) GSVA scores and (C) SCENIC scores of indicated terms or transcription factors in each cluster of proximal tubular cells from BPTES scRNA-seq datasets.

**Figure S6.**
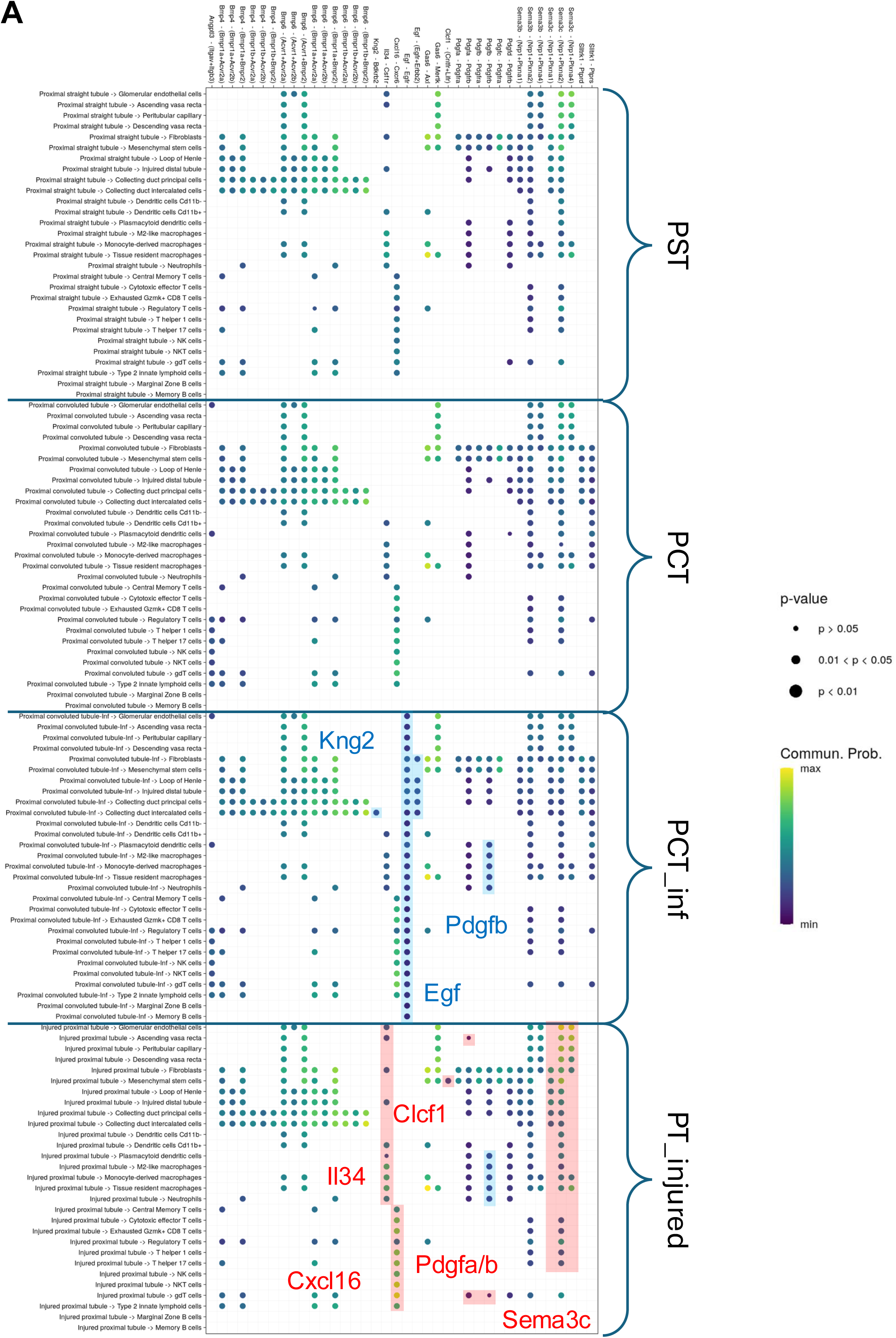
Cell-cell communication inference showed the impact of injured-PT and inflammatory PCT cells which were sensitive to BPTES. The dot plot displays all significantly enriched ligand-receptor pairs from proximal tubular cells to other subtypes. The color of each dot represents the communication probability. Blue rectangles highlight ligand-receptor pairs specific to PCT_inf (C0), while red rectangles indicate those unique to PT_injured.

**Table S1.**
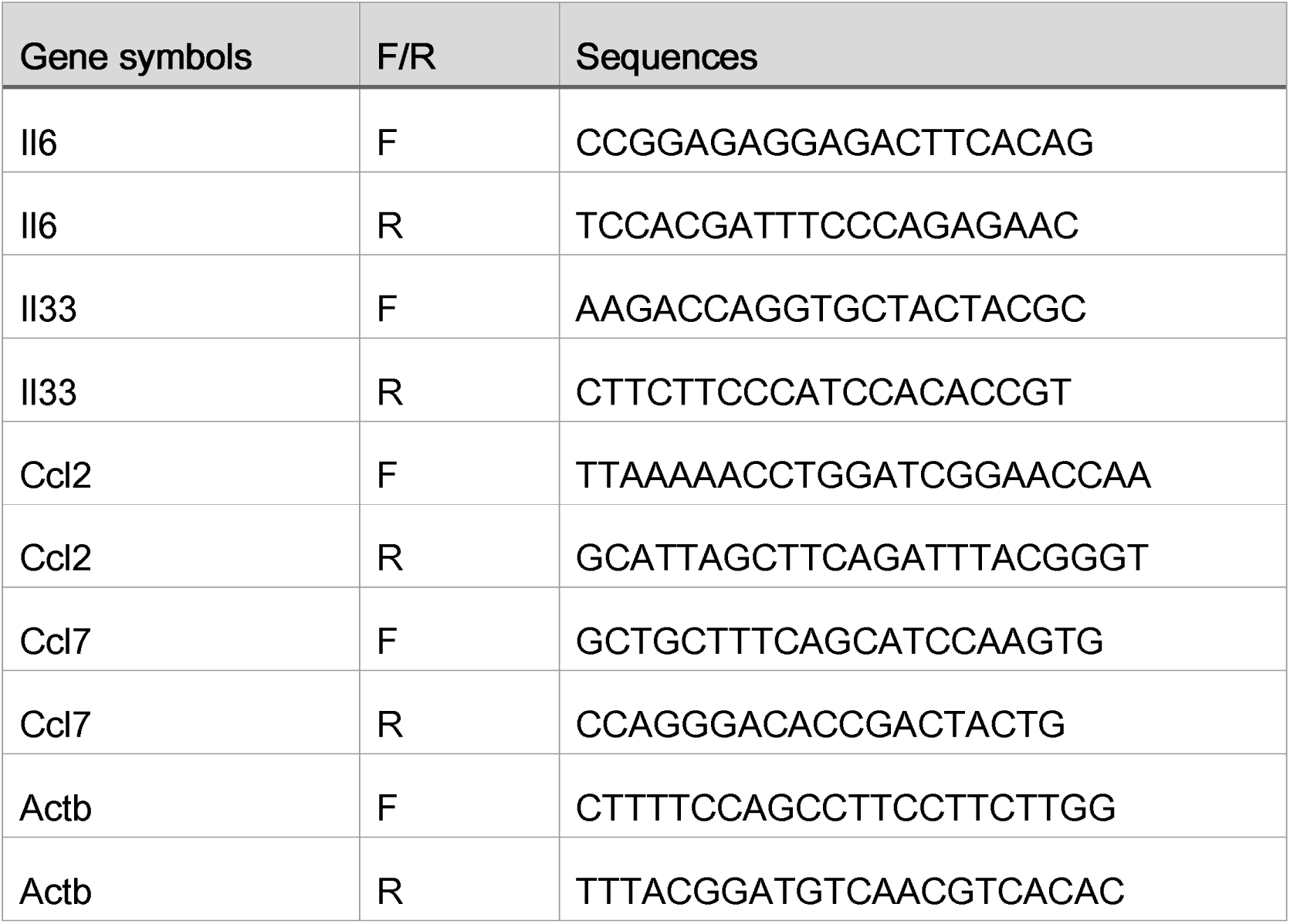
List of qPCR primers

